# Soil microbial responses to multiple global change factors as assessed by metagenomics

**DOI:** 10.1101/2024.07.05.602153

**Authors:** Álvaro Rodríguez del Río, Matthias C. Rillig

## Abstract

Anthropogenic activities impose multiple concurrent pressures on soil ecosystems at a global scale, but the response of soil microbes to multiple concurrent global change (GC) factors is poorly understood. Here, we applied 10 GC treatments individually and in random combinations of 8 factors to soil samples, and monitored their bacterial and viral composition by metagenomic analysis. The application of multiple GC factors selects for particular prokaryotic and viral communities different from the effects of any individual factor, favoring, for instance, potentially pathogenic unknown mycobacteria and novel viruses. At the functional level, multiple GC factors select for sessile and non-biofilm-forming bacteria which are metabolically diverse and show a high load of antibiotic resistance genes. Finally, we show that novel genes are also relevant for understanding microbial response to GC. Our study indicates that multiple GC factors impose directional selective pressures on soil prokaryotes and viruses not observed at the individual GC factor level, and improves our understanding of how GC interactions shape microbial communities.

## INTRODUCTION

Human pressures are numerous, highly diverse in nature ^1^, and influence soil ecosystems at a global scale. One of the most important effects of global change (GC) are shifts in soil microbial populations, central to soil functioning. Several experiments have revealed the response of soil biota to alternative GC factors like warming ^2^, drought ^3^ or microplastics ^4^, among others ^5^. These microbial disturbances impact important soil traits, and monitoring them remains relevant for understanding anthropogenic impacts on soil ecosystems.

However, most studies only include a limited number of GC factors, even though many may act concurrently in natural conditions. In order to address this gap, Rillig *et al.*(2019) ^6^ designed a multifactor experiment including 10 GC factors of diverse nature^1^: warming (physical factor), drought, nitrogen deposition, increased salinity and heavy metals (inorganic chemical factors), microplastics (particle contamination) and antibiotics, fungicides, herbicides and insecticides (organic chemical toxicants). After applying them individually and in random combinations of 2, 5, 8 and 10 to soil samples, results showed that multiple concurrent GC factors triggered directional shifts in soil properties. For instance, individual GC factors barely affected water drop penetration time, but the application of multiple concurrent factors caused a significant increase, more pronounced as the number of applied factors increased. These results highlighted the importance of studying not only the effect of individual GC factors, but also the combined effect of many. Increasing GC factors also triggered directional changes on soil fungal populations, but whether other microorganisms follow the same patterns remains unknown. These include prokaryotes, which are central to soil functioning and usually show different dynamics than fungi ^7,8^, and viruses, important and understudied players of soil functioning ^9^.

In order to understand the response of prokaryotes and viruses to multiple GC factors, we leveraged 70 samples from the multi-factor experiment by Rillig *et al* (2019)^6^, including 10 controls, 50 single GC factor samples (5 samples treated with each individual factor), and 10 multiple GC factor samples (treated with 8 random concurrent GC factors), and analyzed them following a comprehensive metagenomic exploration.

Soils are the most biodiverse habitat on the planet ^10^, and contain an immense number of uncultivated microbial species not present in reference databases ^11^. These unknown species can be uncovered by constructing Metagenome-Assembled Genomes (MAGs) *de novo*, a method that is revealing a great degree of unknown biodiversity ^12^. We recovered a total of 742 mostly unknown bacterial and 1,865 viral MAGs from the 70 soil samples, and leveraged them to describe microbial populations under different GC conditions. We also built a 25M gene catalog to monitor changes at the gene functional level, analyzing both known and novel genes. Our analyses indicate that multiple GC factors have a distinctive effect on soil prokaryotic and viral populations and in the distribution of microbial genes.

## RESULTS

### A genome resolved bacterial catalog

After comparing the performance of different prokaryotic binning strategies (Table S1), we restricted our analysis to the genomic bins computed with the multi-sample binning strategy by SemiBin2 ^13^, which provided 653 medium quality (MQ, completeness >= 50%, contamination < 10%) and 89 high quality (HQ, completeness >= 90%, contamination < 5%) genomic bins^14^. These MAGs show variable genome sizes across treatments: genomic bins reconstructed from samples treated with heavy metals, salinity and the random combination of 8 GC factors are significantly larger than in the control samples (Two-sided Wilcoxon test p-value < 0.01, Figure 1C).

**Figure 1.**
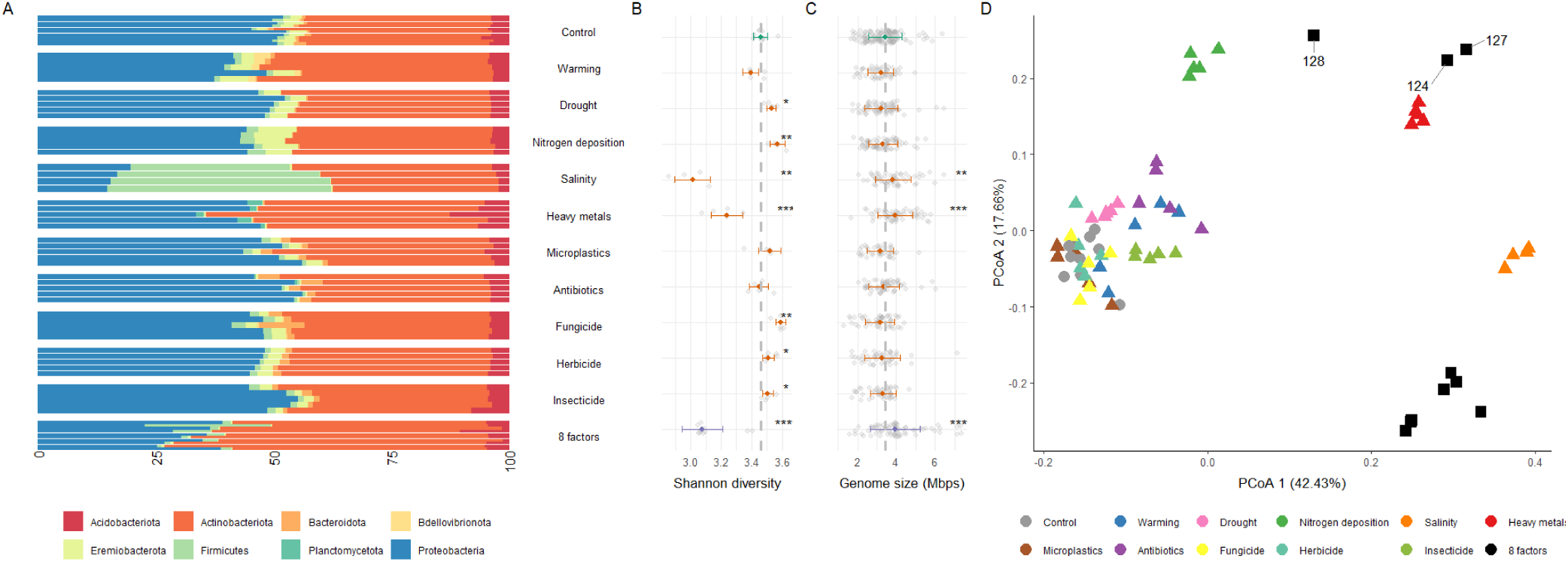
Bacterial composition and diversity change across GC conditions. A) Taxonomic profile of the representative MAGs reconstructed from the 70 samples included in this analysis (10 controls, 5 for 10 different GC factors and 10 random combinations of 8 GC factors) collapsed to the phylum level; B) Shannon diversity index, per treatment; C) MAG genome size, per treatment; D) Principal Coordinate Analysis (PCoA) on the relative abundance of the reference MAGs. Shapes indicate the number of GC factors (Circle = 0, triangle = 1, square = 8). In B and C, thick points represent the median values, and black bars indicate standard deviation intervals. Asterisks represent different significance levels obtained after a Two-sided Wilcoxon test with control samples; * indicate p-value <= 0.05, ** p-value <= 0.01, *** p-value <= 0.001 and **** p-value <= 0.0001.

MAGs across samples were highly redundant and clustered into 77 reference species bins according to the 95% Average Nucleotide Identity (ANI) species definition. Even though we could resolve the phylum, class and order of all, 96.1% reference MAGs could not be taxonomically classified to the species level with GTDB-tk2^15^, indicating they represent unknown taxa. For simplicity, we refer to bins not assigned species-level taxonomic labels as unknown.

### Different bacterial diversity and composition under alternative GC scenarios

We next exploited the reference genomic bins for understanding the effect of different GC factors to bacterial populations. We observed clear diversity shifts across treatments (Bray-Curtis beta-diversity ANOVA p-value < 0.0001, Figure 1A). For instance, most GC scenarios, except antibiotics, drought, warming and fungicides significantly reduced the relative abundance (Two-sided Wilcoxon test p-value < 0.05) of a potentially nitrogen-fixing unknown MAG within the Bradyrhizobium genus, ubiquitous and globally abundant in soil ^16^.

Most individual global change factors lead to significant differences in bacterial alpha diversity compared to control samples (Figure 1B). For instance, heavy metals and salinity trigger significant diversity losses and drive the strongest responses (34.7 and 28.5% mean decrease accuracy in a random forest regression, respectively). In contrast, other factors such as fungicide, nitrogen deposition and drought significantly increased alpha diversity. However, the random combinations of 8 GC factors, regardless of their identity, always decrease bacterial diversity, and the number of factors have a more important contribution in explaining diversity patterns than most individual treatments (9.5% mean decrease accuracy, 5th ranked treatment).

Community composition was markedly different after heavy metal, nitrogen deposition, salinity and the 8 factor treatments (Figure 1D). For instance, salinity samples are characterized by an abrupt increase in the relative abundance of Firmicutes and Proteobacteria bins (Two-sided Wilcoxon p-value < 0.01, Figure 1A). The 8 concurrent factor samples have the highest intra-treatment variability, but also show common community patterns distinct to the control samples (Bray-Curtis beta-diversity ANOVA-like pairwise permutation test p-value < 0.01). For instance, they repeatedly show an increased abundance of Actinomycetia class genomes (Figure 1A), driven by an unknown bin within the Chersky-822 genus, which doubles in relative abundance to become the second most abundant species (Two-sided Wilcoxon p-value < 0.01). Along the first PCoA axis, 8 factor samples cluster together with copper and salinity samples, indicating similarities in their composition. However, they distribute differently across the second axis and form two well differentiated clusters. The first is composed of 7 samples which include the salinity treatment, and the second consists of the 3 remaining samples, which miss the salinity treatment (sample numbers 124,127 and 128). The biggest difference between these two clusters was the abundance of Actinobacteriota genomes, which increased in samples 124, 127 and 128 (Two-sided Wilcoxon p-value < 0.01) while decreasing in the remaining 8 factor samples (Two-sided Wilcoxon p-value < 0.1). The 8 GC factor samples treated with salinity also exhibit a reduced abundance of Proteobacteria (Two-sided Wilcoxon p-value < 0.001), similarly to the individual salinity treatment, which was not observed in samples 124, 127 and 128.

The 77 representative MAGs captured less than 10% of the metagenomic reads, and miss some ubiquitous and abundant soil members such as Rhodoplanes ^16^, indicating they represent an incomplete picture of biodiversity. This is typical of *de novo* genome building ^17,18^, especially in complex environments like soil ^12,19^. Also, given that MAGs were directly reconstructed from samples exposed to the different treatments, the changes observed could arise because of a bias in the reference sequences used. Hence, we confirmed the community patterns on taxonomic profiles built on external reference databases. As many reference-based strategies with different strengths and limitations exist ^19^, we tested different taxonomic profiling methods. Alternative approaches detected a variable number of species, but they all agree in their bacterial composition measurements. In contrast, alpha diversity was more variable, especially after predictions based on read taxonomic classification (Figure S1).

### Multiple global change factors drive an increase of mycobacteria

Some individual factors like salinity, heavy metals, drought and nitrogen deposition increased the relative abundance of 5 reference bins from unknown species within the Mycobacterium genus (Wilcoxon test p-value < 0.05). This increase was especially evident after the application of 8 concurrent GC factors (Two-sided Wilcoxon test p-value < 1e-4, Figure 2A), where the mycobacterium MAGs are among the 12 most abundant genomes, whereas in the control samples the most abundant is ranked 27th. Many Mycobacterium species, known as Non Tuberculous Mycobacterium (NTM) are environmental opportunistic pathogens ^20^, and are becoming an increasing sanitary problem ^21^. In order to understand the potential pathogenicity of the unknown mycobacterium MAGs thriving in multiple GC conditions, we examined their virulence factor content ^22^, which, despite not being markers for pathogenicity, contribute to the ability of pathogens to survive within hosts ^23^. All unknown mycobacterium bins contained a high content of virulence factor genes compared to the rest of MAGs (Figure S2) and similar to other soil NTM MAGs reconstructed by Bin Ma *et al.* (2023) ^24^ (Figure S3).

**Figure 2.**
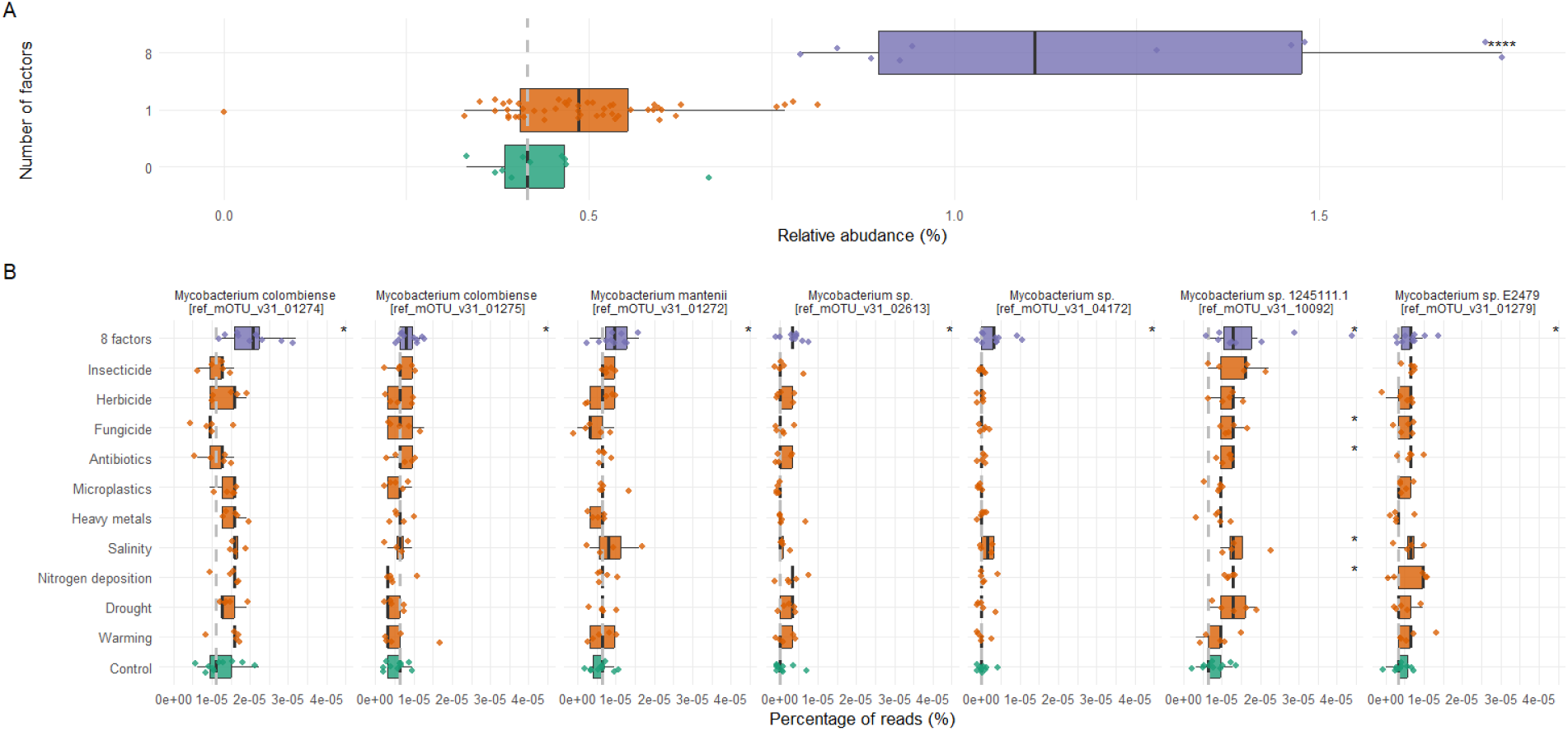
Mycobacterium genomic bins are enriched in multiple GC factor samples. A) Relative abundance of mycobacterium bins in control, one factor and 8 factor samples; B) Abundance of the 7 mycobacterium OTUs significantly enriched after the 8 factor treatment (Two-sided Wilcoxon test p-value < 0.05). Data are represented as boxplots in which the middle line is the median, the lower and upper hinges correspond to the first and third quartiles, the upper whisker extends from the hinge to the highest value no further than 1.5 × interquartile range (IQR) from the hinge and the lower whisker extends from the hinge to the lowest value no further than 1.5 × IQR of the hinge. Asterisks represent different significance levels obtained after a Two-sided Wilcoxon test with control samples; * indicate p-value <= 0.05, ** p-value <= 0.01, *** p-value <= 0.001 and **** p-value <= 0.0001.

Among them, the five unknown reference bins harbor virulence factors characteristic of *Mycobacterium tuberculosis*, which share genomic characteristics with NTM ^25^. For instance, they all show copies of *MmpL13*, essential for the integrity of the mycobacterial envelope ^26^ and type VII secretion systems, which transports proteins across this outer membrane and play a significant role in mycobacterial virulence ^27^. Four bins also contain *Lsr2* genes, involved in the metabolism of *Mycobacterium tuberculosis* during chronic infection ^28^. Additionally, one MAG carries a phospholipase C, which contributes to phagosome escape and facilitates spread ^29^, and three contain copies of *MgtC*, a membrane protein that promotes survival of intracellular pathogens ^30^.

Only 5% of the genes classified as mycobacterium were included in mycobacterium bins, indicating that MAGs did not capture the whole genus diversity. Hence, we next asked whether other already described mycobacterium species may also increase in abundance after the application of multiple GC factors. For this purpose, we exploited the taxonomic profiles built with mOTUs ^31^, which detected 55 Mycobacterium Operational Taxonomic Units (OTUs). Seven OTUs significantly increased in abundance in the 8 GC factor samples (Two-sided Wilcoxon test p-value < 0.05), including *Mycobacterium colombiense* and *Mycobacterium manitenii,* both of which already caused human infections^32,33^. (Figure 2B).

To our knowledge, NTM infections from soil have not been reported. In contrast, fresh water is considered to be the main source of NMT human infections ^34^. Using data from a previous study by Romero et al. (2020) ^35^, we tested whether multiple GC factors (warming and pesticides) increased the relative abundance of mycobacterium in river biofilms. Even though we could not assign any mycobacterium Amplicon Sequence Variant (ASV) to the species level, the only mycobacterium ASV showing variability across treatments is significantly enriched after the two factors were applied concurrently (Two-sided Wilcoxon test p-value < 0.05, Figure S4).

### Multiple global change factors select for members of the rare biosphere

Even though they come from the same soil location, 49 reference bacterial bins (63.6%) do not contain any genome reconstructed from the control samples and only became evident after applying some treatments, indicating the power of treatment application for uncovering biodiversity ^36^. Among them, some representative MAGs are not detected in most control samples (median relative abundance = 0), but recurrently show a higher relative abundance after some GC treatments.

For instance, 5 Firmicutes genomes under the detection threshold in control samples gather more than 0.1% of the metagenomic reads after the salinity treatment (Two-sided Wilcoxon test p-value < 0.05). Among them, we found three unknown genus genomes within the Sporolactobacillaceae family, previously reported to have salt - tolerant members ^37^, one of which becomes the most abundant species. Additionally, two conditionally rare unknown bins are classified within the extremophile Alicyclobacillaceae family ^38^. Similarly, 4 genomes mostly undetected in the control samples have relative abundances higher than 0.1% after heavy metal treatment (Two-sided Wilcoxon test p-value < 0.05), including two unknown MAGs from the Edaphobacter genus (within the extremophile Acidobacteriae class ^39^), an unknown genus MAG within the Isosphaeraceae family, and an unknown 17J80-11 genus bin from the Caulobacteraceae family, which has some copper resistant members ^40^. Nitrogen deposition also favors different rare genomes across the bacterial phylogeny (Figure S5).

Conditionally rare taxa are important for environmental responses to perturbations ^41,42^ because they serve as reservoirs of genetic diversity not necessarily encoded by more abundant taxa ^43^, which allow them perform better in determined conditions, and contribute to environmental resistance and resilience ^43–45^. After a differential frequency analysis, we found that the 5 conditionally rare Firmutes genomes enriched after the salinity treatment show a higher frequency of many genes involved in sporulation (q-value < 1e-10, Table S2). Similarly, the four rare genomes enhanced by heavy metals are enriched (q-value < 0.01) in several metal transporter and succinoglycan biosynthesis sequences, which can confer resistance to extreme conditions ^46^, among other genes (Table S3).

### Different global change conditions increase the abundance of unknown viruses

We next asked whether GC conditions also select for different viral populations. For monitoring how viral abundance shifts under different GC treatments, we first identified viral contigs with VirFinder ^47^, and phage contigs with Seeker ^48^. Both viruses and phages increase in frequency after some treatments, especially salinity, nitrogen deposition and the random combination of 8 GC factors (Two-sided Wilcoxon test p-value < 0.0001, Figure S6).

In order to understand which phage species drive these responses, we computed *de novo* phage bins with PHAMB ^49^. We reconstructed 882, 931 and 52 medium quality, high quality and complete MAGs respectively, which de-replicated into 895 reference viral bins. 48.5**%** of them, including 4 complete bins, do not match any reference sequence, and increase in abundance after most treatments, especially the 8 concurrent GC factors (Two-sided Wilcoxon test p-value < 0.0001, Figure 3A). In contrast to bacteria, viral diversity increases after most treatments (Figure 3B), with salinity, warming, heavy metals and antibiotics as the most important diversity determinant (mean decrease accuracy in a random forest regression higher than 18%). However, compositional analyses indicate a differential viral composition under multiple GC conditions, with the 8 factor samples gathering into two well differentiated clusters (Figure 3C), mirroring bacterial composition.

**Figure 3.**
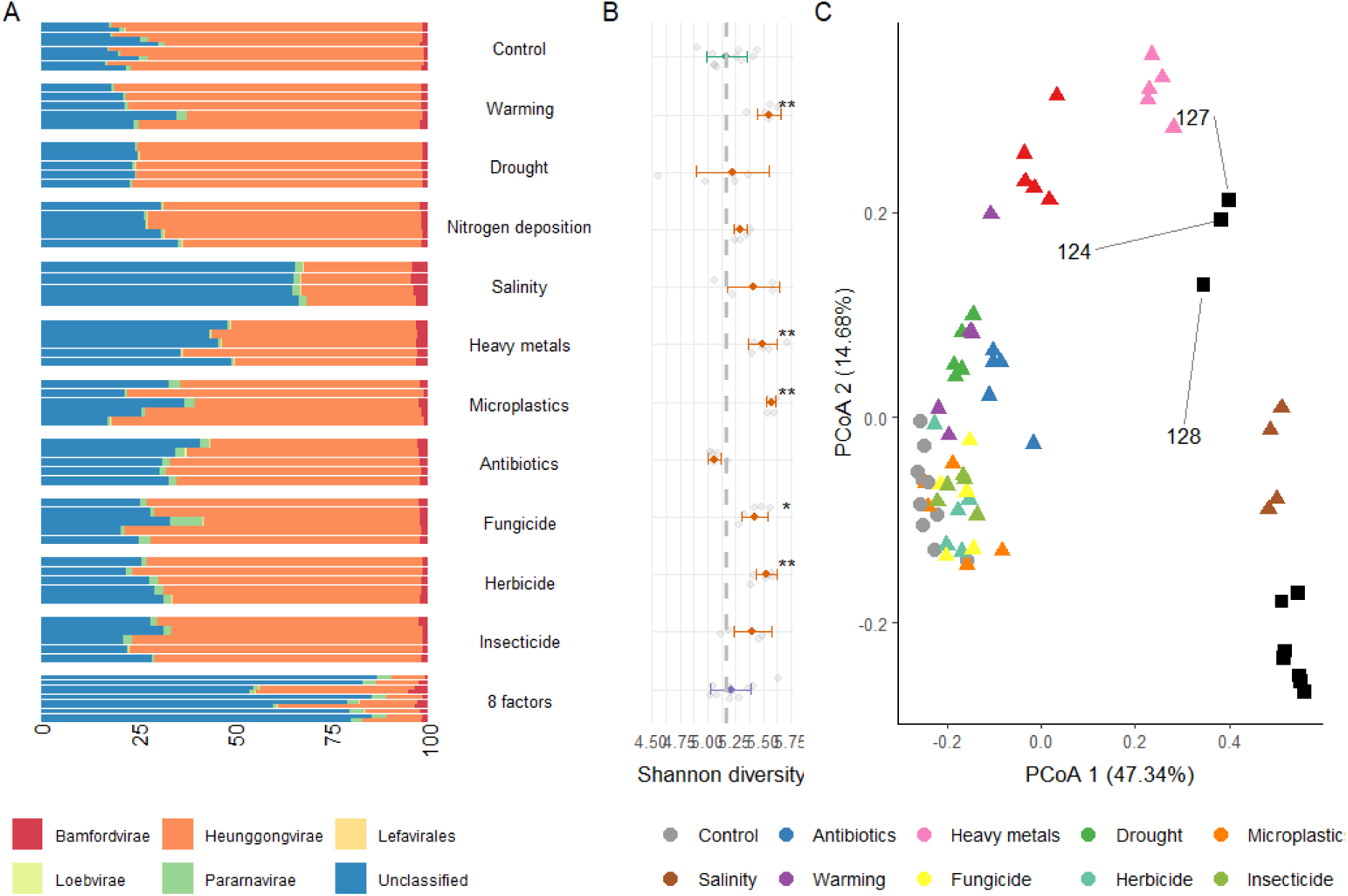
Viral composition and diversity change across GC conditions. A) Taxonomic profile of the representative viral MAGs reconstructed from the 70 samples included in this analysis (10 controls, 5 for 10 different GC factors, 10 random combinations of 8 factors) collapsed to the class level; B) Shannon diversity index, per treatment. Thick points represent the median values, and black bars indicate standard deviation intervals. Asterisks represent different significance levels obtained after a Two-sided Wilcoxon test with control samples; * indicate p-value <= 0.05, ** p-value <= 0.01, *** p-value <= 0.001 and **** p-value <= 0.0001 C) Principal coordinate analysis based on the taxonomic annotations of the reference viral MAGs. Shapes indicate the number of GC factors (Circle = 0, triangle = 1, square = 8).

### Different life history traits under global change

Distantly related species can show similar genes and perform overlapping ecosystem functions ^50^. Hence, we next asked whether the differences observed at the taxonomic level translated into different functional repertoires. For this purpose, we constructed a gene catalog, a strategy widely exploited for describing the functional potential of microbiomes ^51^, including both binned and non-binned contigs. We followed a comprehensive gene prediction strategy for accurately predicting both prokaryotic and eukaryotic genes (see methods), and gathered a total of 25,162,374 genes. The functional profile of samples from the salinity, heavy metal and 8 factor treatments were similar at general functional categories, and formed a separate cluster from the remaining treatments (Figure S7). However, KEGG orthology (KO) composition of the 8 factor samples was mostly different from the rest, including salinity (Figure 4A). For instance, 10 spore germination KOs are significantly depleted in the 8 factor samples, but significantly enriched in the salinity samples (Two-sided Wilcoxon test p-value < 0.05).

**Figure 4.**
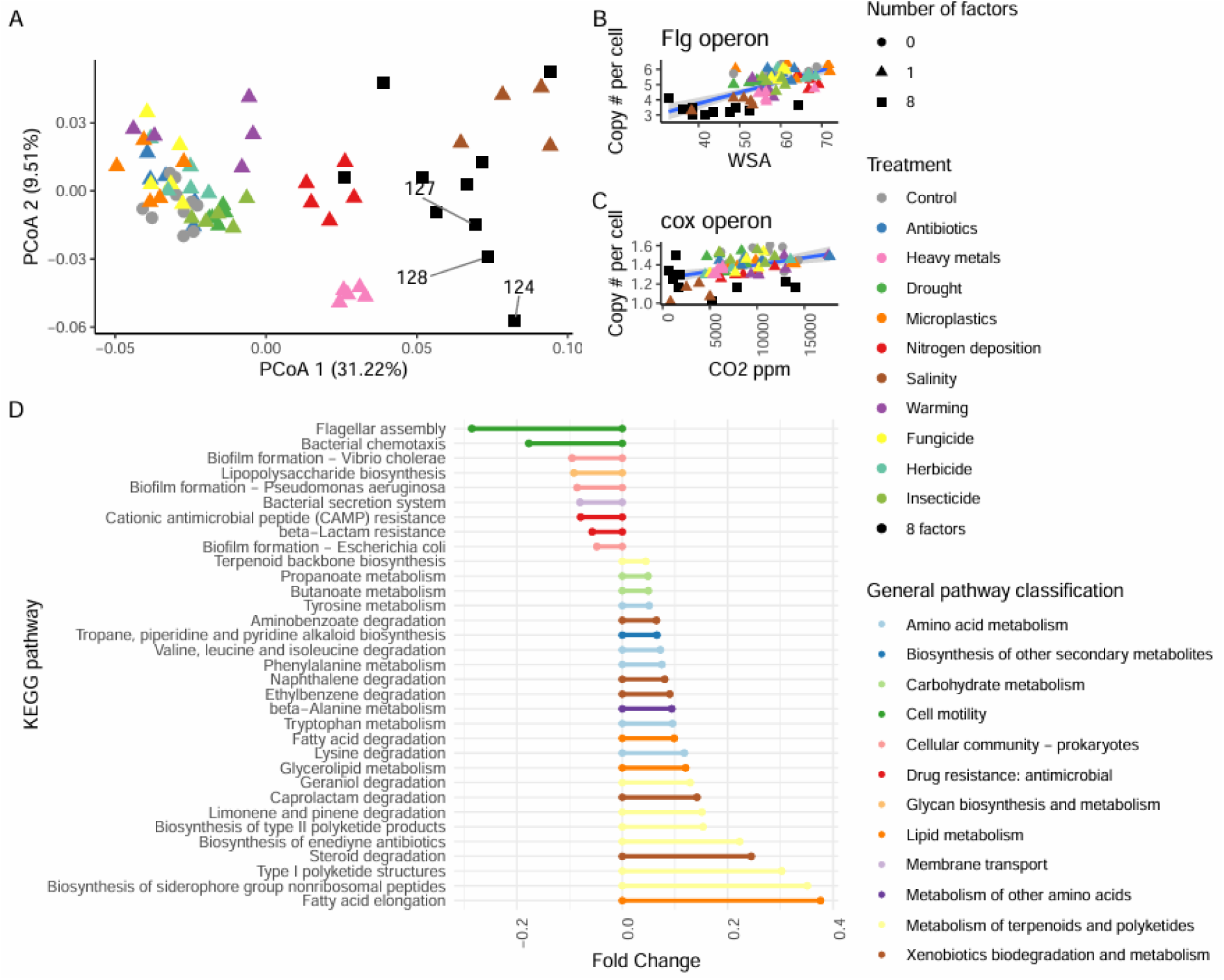
Multiple GC selects for microbial populations with particular gene repertoires A) PCoA built on the frequency of KEGG Orthologues (KOs). We indicate samples 124, 127 and 128, missing the salinity treatment B) Correlation between the frequency of flagellum assembly genes (flg operon) and water stable aggregates (WSA). C) Correlation between the frequency of respiration genes (coxABCD) and soil respiration (CO^2^ ppm). D) KEGG pathways with significant frequency shifts after the 8 factor treatment (q-value < 0.01).

Motility and biofilm formation are considered escape mechanisms for some stress conditions ^52^, but genes related to these processes are significantly depleted when the 8 factors were applied concurrently (q-value < 0.01, Figure 4D). In contrast, these samples show higher frequency of genes involved in several metabolism and degradation pathways. These gene content patterns suggest that the 8 GC factors selected for a nutrient recycling life history strategy characterized by increased assimilation and degradation capabilities, instead of an environmental responsiveness strategy ^53^. An exception were cytochrome oxidase genes, last enzymatic complexes of most aerobic respiratory chains ^54^ and markers for respiration, which are significantly depleted after the 8 factor treatment (Two-sided Wilcoxon test p-value < 0.001) and significantly correlate (R = 0.42, p-value < 0.001, Figure 4C) with CO^2^ measurements. We explored additional associations between soil properties and gene frequencies, and found significant correlations between Water Stable Aggregates (WSA) and genes previously related to soil aggregation (Figure S8). Interestingly, flagellar genes also correlate with WSA (R = 0.59, p-value < 1e-7, Figure 4B), suggesting a relation between bacterial motility and soil aggregation.

### Bacterial composition drives the increase of ARGs in the multifactor samples

Soil is acknowledged to be an important reservoir of Antimicrobial Resistance Genes (ARGs), some of which are or may become clinically relevant ^55^. Hence, it is specially relevant to monitor ARG distribution across GC treatments. In order to understand changes in ARG frequency under different GC conditions, we mapped the genes predicted on each sample against the CARD database ^56^. Despite any individual GC factor, the 8 concurrent factors significantly increased both antibiotic inactivation and antibiotic target protection copy number per cell (Two-sided Wilcoxon test p-value < 0.01, Figure 5A).

**Figure 5.**
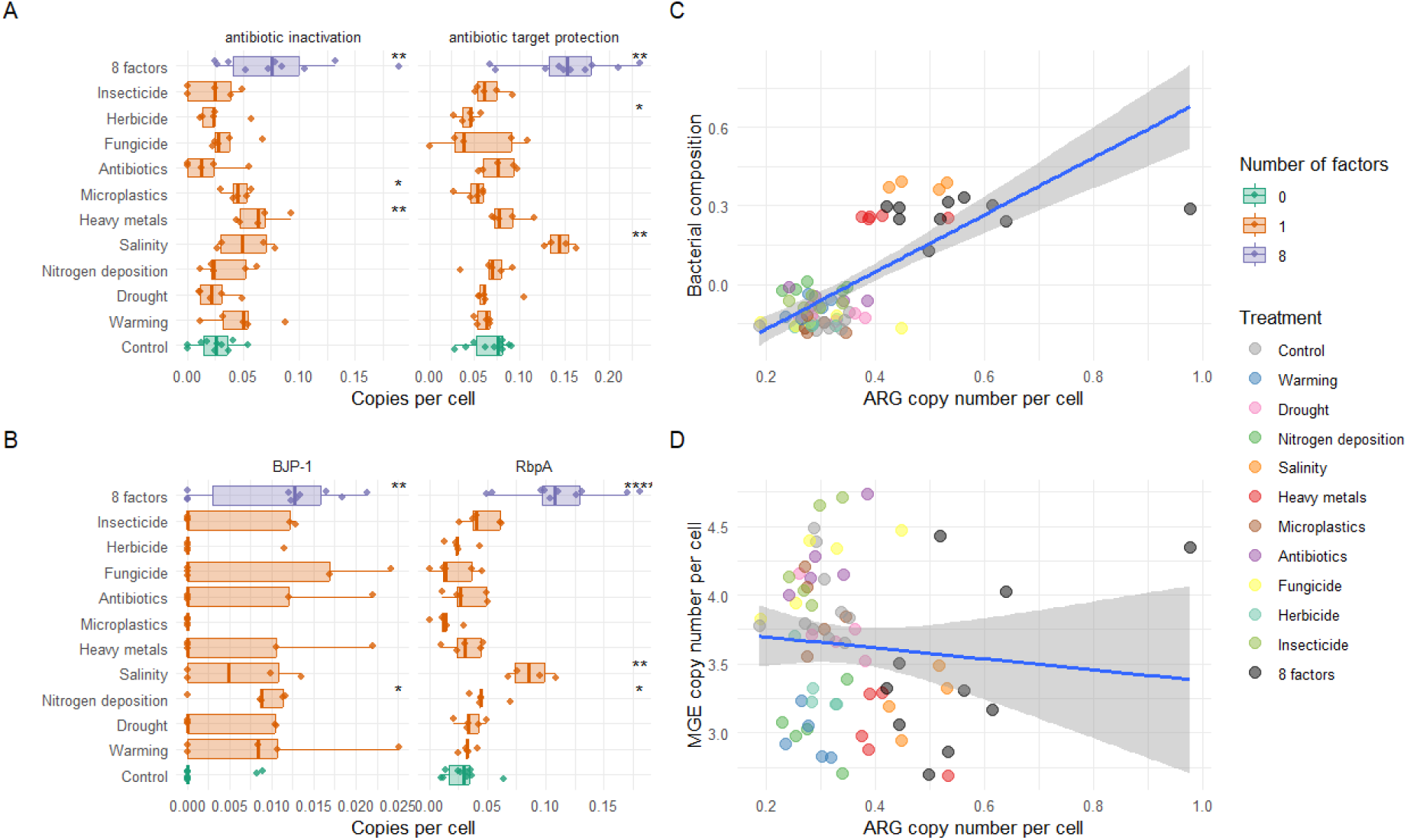
Multiple GC factors drive the increase in abundance of different Antibiotic Resistance Genes (ARGs) A) Antibiotic inactivation and target protection copy number per cell across treatments; B) Copy number per cell variation of BJP1 (antibiotic inactivation) and RbpA (antibiotic target protection) C) Correlation between bacterial composition and ARG copy number per cell; D) Correlation between MGE copy number per cell and ARG copy number per cell. In A and B, data are represented as boxplots in which the middle line is the median, the lower and upper hinges correspond to the first and third quartiles, the upper whisker extends from the hinge to the highest value no further than 1.5 × interquartile range (IQR) from the hinge and the lower whisker extends from the hinge to the lowest value no further than 1.5 × IQR of the hinge. Asterisks represent different significance levels obtained after a Two-sided Wilcoxon test with control samples; * indicate p-value <= 0.05, ** p-value <= 0.01, *** p-value <= 0.001 and **** p-value <= 0.0001.

We did not find a strong correlation between the copy number of ARGs and the total number of mobile genes per cell (R = -0,09, p-value = 0.46, Figure 5D). In fact, only 0.4% ARGs were encoded in contigs classified as plasmids, whereas 1.1% of the genes in the catalog are. Instead, we observed that salinity, heavy metals and 8 factor samples contain higher doses of ARGs, driving a correlation between ARG copy number per cell and community composition (R = 0.56, p-value < 1e-6, Figure 5C) and indicating that phylogeny may be an important driver of ARG frequency, as previously observed in soil ^57^.

In fact, the increased frequency of ARGs in the 8 factor samples was in part driven by the increase in abundance of mycobacterium species. For instance, the antibiotic target protection increase in frequency was mainly driven by the increase in abundance of the *rbpA* gene (Two-sided Wilcoxon test p-value < 0.001, Figure 5B), a RNA-polymerase binding protein which confers resistance to rifampin in mycobacterium ^58^. *EfpA*, a MFS transporter typically found in *Mycobacterium tuberculosis* that confers resistance to several antibiotics ^59,60^, also showed increased frequency after the 8 factor treatment (Two-sided Wilcoxon test p-value < 0.01). Similarly, the only inactivation gene significantly increasing in frequency after the 8 factor treatment was the metallo-beta lactamase BJP-1, characteristic of Bradyrhizobium ^61^ (Two-sided Wilcoxon test p-value < 0.0001, Figure 5B).

### Novel gene families distribute differently across taxa and GC treatments

Because of their unique conditions – e.g. high phylogenetic diversity and number of uncultivated species – soils harbor a large collection of novel genes ^51,62^, which are acknowledged to play important biological roles ^63,64,65^. Hence, we assessed the distribution of novel genes in microbial populations under different GC conditions. Even though only 14.87% of the genes within the catalog lack homologs in EggNOG^66^, they represent 75% of the gene families built by *de novo* after de novo clustering ^67^ (Figure 6A).

**Figure 6.**
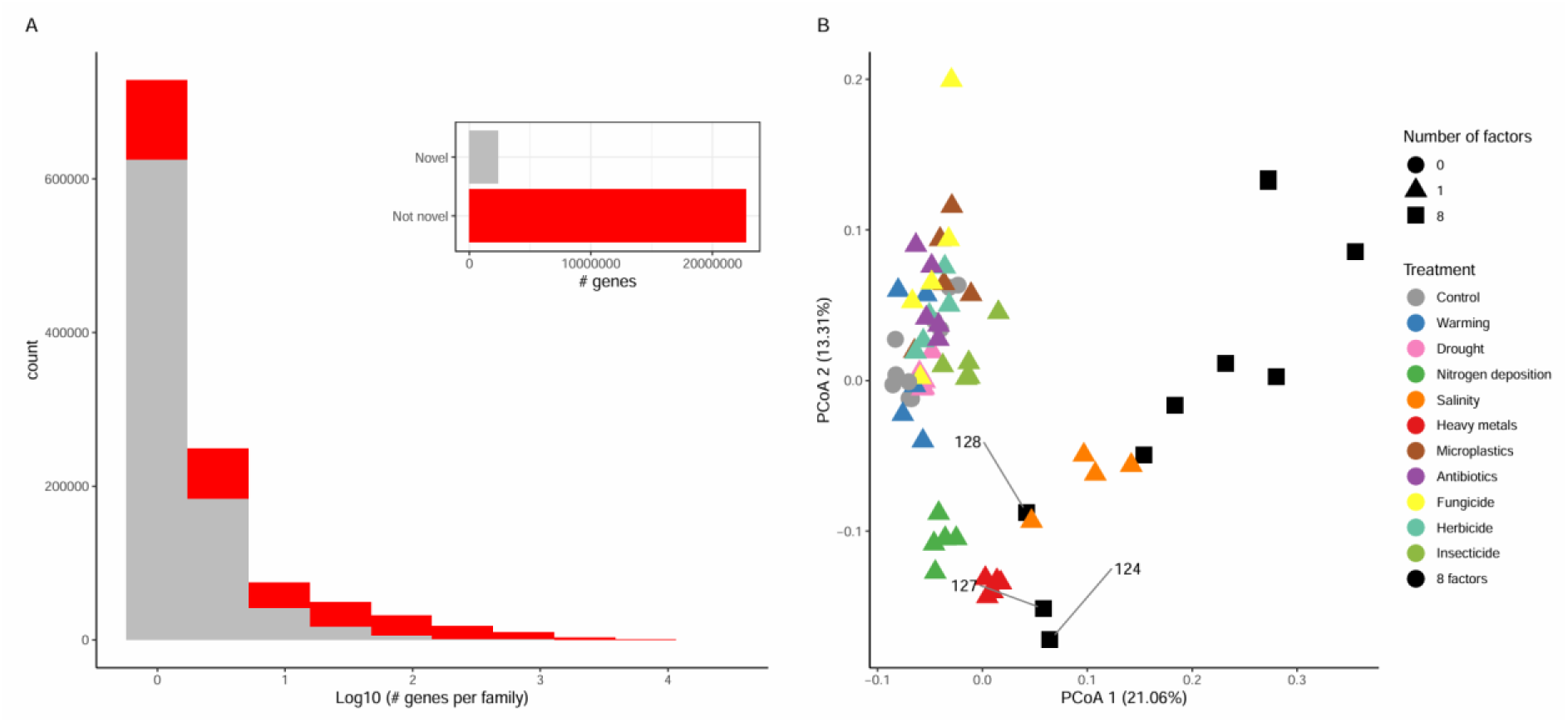
Novel gene families are abundant and distribute differently across GC treatments A) Gene family size distribution. Top right diagram represents the total number of novel (gray) and non novel (red) genes; B) PCoA on the copy number per cell of novel gene families assembled in more than 50 samples. We indicate samples 124, 127 and 128, missing the salinity treatment.

Only 3.9% novel families are encoded in contigs binned into MAGs, but some bins show high degrees of novel gene content. For instance, more than 20% of the genes from unknown Diplorickettsiaceae family bins and Alicyclobacillus genus bins are novel. Additionally, all conditionally rare MAGs contain a higher proportion of novel gene families (e.g. 301 in the genomes enriched after the salinity treatment vs 197 in the rest), some of which may be important to their good performance in the different GC environments. Even though most novel families (71%) are singletons, some show a wide phylogenetic distribution: 1,009 gather more than 100 genes, and 748 are binned in MAGs from different bacterial phyla, classes and orders.

Besides, novel gene families distribute differently across treatments (Figure 6B), and discriminate the 8 factor samples in a coordinate analysis. For instance, 1,163 novel families were exclusively assembled in more than 50% samples from a given treatment, including 64 in antibiotics, possibly representing AMR mechanisms, and 92 in the 8 factor samples. Even though none of these 92 novel gene families were binned into MAGs in our samples, we detected them in other public genomic repositories, where 19 distribute across different bacterial phyla and 5 are specific to mycobacterium (one of them to the NTM *Mycobacterium vulneris* ^68^). Moreover, we found 10 novel gene families significantly overrepresented (q-value < 0.05) in the 8 factor samples, two of them also detected in external mycobacterium MAGs. Describing these families remains critical for understanding microbial populations thriving under different GC conditions.

## DISCUSSION

Here we provide the first report on the effect of multiple concurrent GC factors in soil bacteria and viruses, complementing earlier results on fungal communities and soil properties^6^. Our study includes 10 individual GC factors, most of which have been broadly studied individually, but not within the same soil context. For instance, salinity has large impacts on bacterial communities, as already noted ^69,70^, and selects for conditionally rare taxa that are central for a complete understanding of soil’s response to GC. Despite the strong effect of salinity, we demonstrate that the 8 concurrent factors selected for particular bacterial and viral communities. This is remarkable, as the identity of all the factors included is different ^1^, and every sample applied with 8 random GC factors represents unique conditions which only have the number of factors in common. This outcome is consistent across different methodologies and biodiversity levels tested. However, our analysis focuses on a particular time point (6 weeks after treatment) in a particular soil, and further efforts will be needed to assess whether the strong effect of multiple GC factors is generalizable to other soil types and exposure times.

A distinctive feature of multiple GC samples is the increased relative abundance of potentially pathogenic Mycobacterium genomic bins. Mycobacterium are ubiquitous in soil ^16^ and known for their ability to survive in harsh conditions ^71^, which explains their increased abundance in the highly perturbed multifactor samples. These conditions may mirror urban environments, which show more acute global change than natural environments ^72^, highlighting the possible relevance of this finding to human health. Previous work highlighted that aquatic mycobacterium, more strongly associated with human health, may proliferate under GC conditions ^73^, but the effect on multiple concurrent factors on aquatic mycobacterium abundance is yet unknown. We could confirm that aquatic mycobacterium increases in relative abundance when exposed to two concurrent GC factors, but additional studies will be needed to confirm this pattern with a higher number of GC factors.

Changes at the taxonomic level also translated into shifts at the functional level for most treatments, especially salinity, heavy metals and the 8 concurrent factors, which form a separate cluster when considering general functional categories. However, the 8 factor treatment shows a differential composition of KOs, novel gene families and AMR genes, indicating the different metabolism of the microorganisms thriving under these conditions. Given their increased genome size and enriched frequency of metabolic genes, populations surviving multiple GC factors seem to be metabolically diverse, potentially allowing them to leverage a wider range of compounds. In contrast, motility and biofilm formation, which can be resistance mechanisms ^74,75^, are not selected under multiple GC conditions.

We found associations between the frequency of particular genes and previously reported soil processes. For instance, CO^2^ measurements correlate with the frequency of *cox* genes, central for soil respiration. Similarly, motility genes significantly correlate with WSA, indicating an impact on one another, although in this case the direction of the relation is not straightforward. On the one hand, a meta-analysis suggested that sessile bacteria have a stronger effect on soil aggregation than motile bacteria ^76^. On the other hand, motile bacteria may not perform well in low WSA soils because they need water films to swim ^77^, and such pore spaces may be less optimal in less aggregated soils. Moreover, they may be more exposed to GC factors while moving through soil, decreasing their fitness under multiple GC conditions.

Finally, we also highlight the high abundance of novel gene families in soil samples, and demonstrate that they are relevant for understanding the biology of microbial populations thriving in different GC environments. In fact, even though mostly unbinned, they provide a good discrimination of the 8 factor samples, indicating that uncultivated species not captured by MAGs also show a differential response to multiple GC conditions.

## METHODS

### Experimental design

This study used samples from a controlled environment experiment conducted with 10 factors of global change on soils. Samples from this experiment were immediately frozen at the time of harvest for the analyses described here. For details see Rillig et al. (2019)^6^. Briefly, the experiment used mini-bioreactors with 30g soils to which 10 different GC treatments were applied, either individually, or in combination.

The 10 global change factors considered were i) warming (increment of 5.0°C over an ambient temperature of 16.0°C); ii) Nitrogen enrichment (added the equivalent of 100 kg N ha^-1^ yr^-1^ ammonium nitrate to the experimental units in dissolved form); iii) Drought (added half of the amount of water at the beginning of the experiment, compared to control water levels that were at 60% of water holding capacity); iv) Heavy metal (copper (ii)-sulfate - pentahydrate to the soil in dissolved form to a final concentration of 100 mg Cu kg-1); v) microplastics (polyester fibers at a concentration of 0.1%); vi) Salinity (added NaCl to the soil until 4.0 dS m^-1^); vii) Herbicide (50 mg kg^-1^ of Roundup® PowerFlex (Monsanto Agrar Deutschland, Düsseldorf), which contains 480 g L^-1^ glyphosate as active ingredient); viii) Antibiotics (applied oxytetracycline at concentrations of 3.050 mg kg^-1^); ix) Insecticide (50 ng g^-1^ of imidacloprid) and x) fungicides (6.0 mg kg^-1^ of carbendazim).

For the 8-factor combination, treatments were drawn at random without replacement for each replicate from the set of 10 treatments. This treatment therefore emphasizes the co-occurrence of 8 factors of global change, while de-emphasizing (through the random draws) the composition of factors. The experiment lasted six weeks, a period sufficient for effects to manifest in such soil experimental systems.

### Shotgun sequencing

Genomic DNA was extracted with the Qiagen DNA Power Soil kit on 250 mg of soil after mixing each sample for homogenization. The genomic DNA was randomly sheared into fragments, which were end repaired, A-tailed and further ligated with Illumina adapters. The fragments with adapters were size selected, PCR amplified, and purified. Sequencing of the 150bp paired-end reads was performed on an Illumina Novaseq 6000 platform using V1.5 reagent and a S4 flow cell.

### Read processing

Reads obtained from the shotgun metagenome sequencing of the soil samples were trimmed as follows: i) Adapters were removed; ii) Repetitive/overrepresented sequences generated by FASTQC reports were trimmed; iii) Tandem repeats were discarded with the TRF software ^78^; iv) Reads were cut when the average quality per base drops below 20, after scanning the read with a 4-base wide sliding window; v) Leading and trailing “N” bases or bases with quality lower than 3 were removed.

Reads below 50 bases after trimming, and reads matching to the human genome (hg37dec_v0), were discarded. These filtering steps were run with the kneaddata software (available at https://github.com/biobakery/kneaddata). We rarefied the samples to the library size of the lowest coverage sample using seqtk ^79^. Rarefied sequences were only used for quantification purposes.

### Assembly

Contig assembly was performed with SPAdes ^80^ with the --meta -m 500 --only-assembler parameters. Contigs smaller than 1000 bps were discarded for subsequent analysis.

### Prokaryotic binning

We binned the assembled contigs with: i) SemiBin2 ^13^ with the multi_easy_bin option after mapping the reads to the contigs with BWA ^81^; ii) MaxBin 2.0 ^82^ and iii) Metabat2 2 ^83^(-m 1500 -s 100000 flags). We also merged the predictions of the 3 of them with MAGScoT ^84^ with default parameters, and with more relaxed options (-t 0.2 -m 10 -a 0.5 -b 0.2 -c 0.2). Genome quality was calculated with CheckM2 ^85^. SemiBin2 provided the highest number of bins (Table S1), which were used for subsequent analysis. We ran dRep ^86^ for deduplicating the bins and generating representative species bins (95% ANI threshold). We then assigned taxonomic labels to the bins with gtdbtk-2.1.0 ^15^, using the r207 GTDB database as a reference. The relative abundance of each representative bin on each sample was calculated by mapping the rarefied reads against the genomes’ contigs with CoverM (available at https://github.com/wwood/CoverM, --min-read-percent-identity 95 --min-read-aligned-percent 75 -- methods relative_abundance).

For running the phylogenetic tree of the reference MAGs, we first identified marker genes. For that purpose, we mapped the gene predictions of the bins (run with prodigal, see gene prediction section) against the marker genes from the GTDB_r214 ^87^ database. Hits with an e-value < 1e-3 were considered as significant. We then computed gene alignments for each marker gene with MAFFT ^88^ (--localpair --maxiterate 1000 options), and discarded position with gappiness > 80% with trimAl ^89^. The final tree was run with IQ-TREE ^90^ -nt AUTO -m GTR+G -cptime 5000 options. For identifying genomes enriched in particular KOs, we used the mannwhitneyu python function, and applied the Bonferroni method for correcting for multiple testing.

### Contig classification

We classified contigs into plasmid / chromosomal with i) Plasflow ^91^ (default options), ii) PlasmidHunter ^92^ (default parameters) and iii) RFPlasmid ^93^ (--species Generic and --jelly flags). Contigs predicted as plasmids by the three software were considered to be so. For locating viral contigs, we used VirFinder ^47^, and considered as viral those contigs with p-value < 0.05. For locating phages, we ran Seeker ^48^ on the viral contigs (predict-metagenome script, default options). We also ran Whokaryote ^94^ with default options to identify eukaryotic contigs.

### Viral binning

Viral bins were calculated using the PHAMB software ^49^, on the bins calculated with VAMB ^95^. Viral bin quality was assessed with CheckV ^96^, and bins labeled as medium quality, high quality and complete were dereplicated with dRep (95% ANI). Viral taxonomy was obtained from the geNomad taxonomy^97^ provided by CheckV.

### Gene prediction

For running gene predictions, we ran MetaEuk ^98^ (easy-predict flag) for the contigs classified as eukaryotic by Whokaryote, and Prodigal ^99^ (-p meta and -f gff parameters) for the contigs classified as non-eukaryotic. We predicted a low number of genes in eukaryotic contigs (337,262, 1.3% of the total number of genes), as also found in other studies ^100^. Eggnog-mapper v2 ^101^ was run for obtaining the functional annotation of the genes (--itype proteins --block_size 0.4 options).

We also identified genes within the CARD ^56^, mobileOG-db ^102^ and VFDB ^22^ databases by mapping the protein sequences with DIAMOND ^103^ blastp and the sensitive flag. Hits with an e-value < 1e^-7^, similarity > 80% and coverage > 75% were considered as significant ^104^. We followed the same approach for locating genes from the VFDB database in public soil MAGs^24^.

Flagellar genes within the *flg* operon considered were K02481, K02482, K02386, K02387, K02388, K02389, K02390, K02391. K02392, K02393, K02394, K02395, K02396, K02397, K02398, K02399 (*flgABCDEFGHIJKLMNRS*). Genes within the cytochrome c oxidase cox operon were K02274, K02275, K02276, K02277 (*coxABCD*). WSA related genes considered were K01991 (polysaccharide biosynthesis/export protein, *gfcE*), K09688, K09689 and K10107 (capsular polysaccharide export systems *KpsMTE*), K07091, K09774, K11719, K11720 (lipopolysaccharide export proteins *LptFACG*) and K04077 (H*SP60/GroEL*)^105^.

### Gene frequency quantification

Copy number per cell was recently recommended for gene quantification in metagenomes ^106,107^. We also decided to use this estimate, instead of relative abundances, because the number of marker genes detected after the different treatments was markedly different (Figure S9). This may be because of a different proportion of eukaryotes or viruses (Figure S6).

We calculated copy number per cell for each KEGG ^108^ Orthology, KEGG pathway, CARD and mobile-OG genes. For doing so, we first calculated the mean number of marker genes per sample by mapping the HMMs of the 41 COGs within the fetchMG database (available at http://motu-tool.org/fetchMG.html) with HMMsearch ^109^ against our gene predictions. Hits with E-value < 1e-3 were considered as significant. We then calculated the copy number per cell of a given annotation X in a sample Y as: Number of genes homologs to X detected in sample Y / Average number of marker genes in sample Y.

For calculating differentially abundant gene pathways, we computed a Wilcoxon test as implemented in the ‘coin’ R package, correcting the p-values by the FDR method to adjust for multiple testing. prokaryotic KEGG pathways with q-value < 0.01 were considered to be differentially present in control and treated samples.

### Reference based taxonomic profiling

Given the different performances of alternative taxonomic profiling methods ^19^, we followed a comprehensive approach, gathering results from different software and reference databases. Taxonomic profiling was performed on the rarefied reads with Kraken2 ^110^, using the k2_pluspf_20230605 database as reference with the --use-names flags. We also identified species within the SILVA ^111^ 138.1_SSURef_NR99_tax_silva_NR97 databases by mapping the rarefied reads with bowtie2 (--very-sensitive flag). We also computed species abundance by identifying marker genes, regarded to be a more accurate approach ^107^ with mOTUs ^31^ v3.1.0 (-t 1 -A -c -q flags) and SingleM ^11^.

### Species DA analysis on a multiple stressor experiment on water

In order to confirm that mycobacterium species increase in abundance because of multiple factors in water, we downloaded the data generated by Romero et al (2020)^35^ from the NCBI (accession number PRJNA574152). We ran the DADA2 pipeline ^112^ for quantifying the relative abundance of mycobacterium species.

### Novel gene family identification and analysis

We clustered all the gene predictions into gene families with MMseqs2 ^67^ relaxed parameters (--min-seq-id 0.3 -c 0.5 --cov-mode 1 --cluster-mode 2 -e 0.001). We considered gene families with no members detected with eggnog-mapper^101^ as novel (i.e. not present in reference species). We located novel families in external MAGs by mapping their longest representative against the proteins encoded in a collection of 169,484 genomes spanning the prokaryotic tree of life^12,51,87,113,114^. We used DIAMOND blastp (‘sensitive’ flag). Hits with an e-value < 1e 10^−3^ and query coverage >50% were considered as significant).

### Statistics and figures

Figures were generated with the ggplot2 R package. Tree figures were generated using the ggtree R package. Heatmaps were generated with the pheatmap R function. Shannon diversity was calculated with the diversity function within the microbiome R package. Beta diversities were calculated with the vegdist function within the vegan database, using Bray-Curtis distances. Permutations of the multivariate homogeneity of group dispersions (variances) were calculated with the betadisper and permutest R functions. PCoAs were built using the ape pcoa function, providing Bray-Curtis distance matrices computed with the vegan vegdist function. The relative importance of each factor was calculated with the randomForest R function. In python,we used the pandas, scipy and numpy libraries. Sample 85 (salinity treatment) was discarded from the statistical analysis because it shows a high deviation compared to other samples.

## ACKNOWLEDGMENTS

ARdR was funded by a Humboldt research fellowship for postdoctoral researchers. We thank Anja Wulf for extracting DNA and Stefan Hempel for organizing the sequencing.

## DATA AVAILABILITY

The sequences from the 70 metagenomic samples have been deposited in the NCBI under Bioproject code PRJNA1102178.

## CODE AVAILABILITY

The code used for the analysis have been deposited in https://github.com/AlvaroRodriguezDelRio/Multiple_GC_soil_experiment/

## Supplementary figures

**Figure S1.**
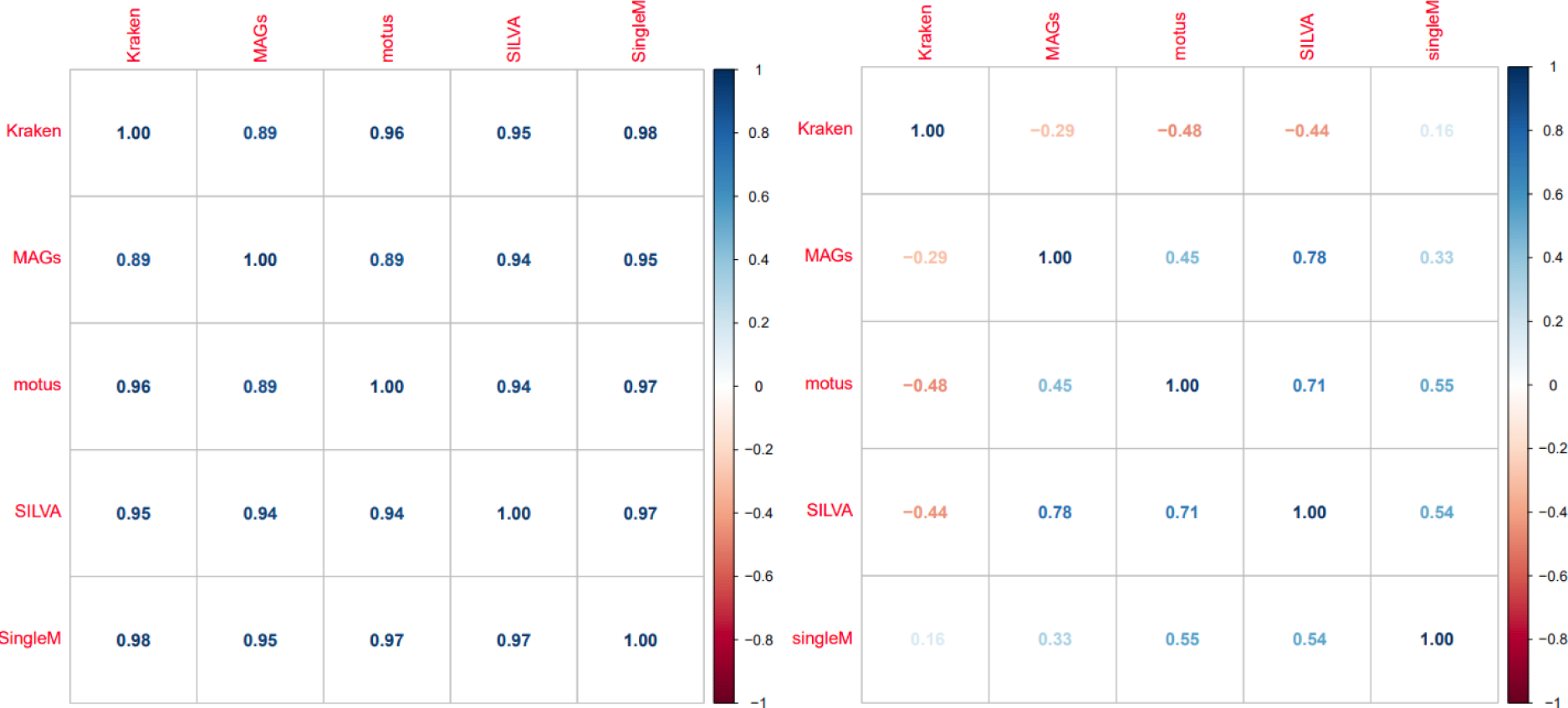
Correlation across bacterial composition (right) and diversity (left) measured by different taxonomic profiling methods. Kraken (read taxonomic classification) provides different diversity patterns than the rest of methods.

**Figure S2.**
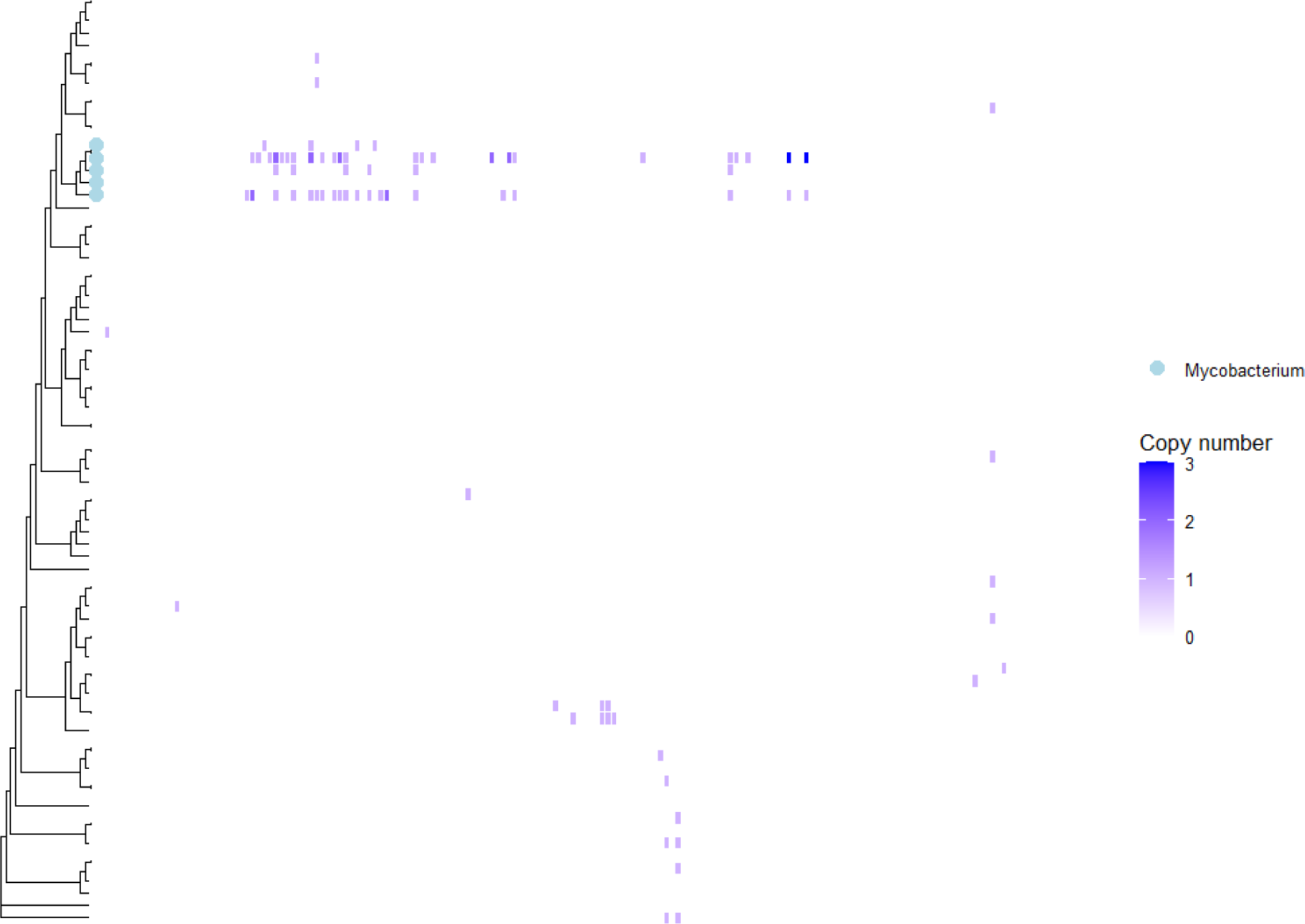
Virulence factor content (columns) per reference bin (rows). Blue tips point to the five Mycobacterium genomes

**Figure S3.**
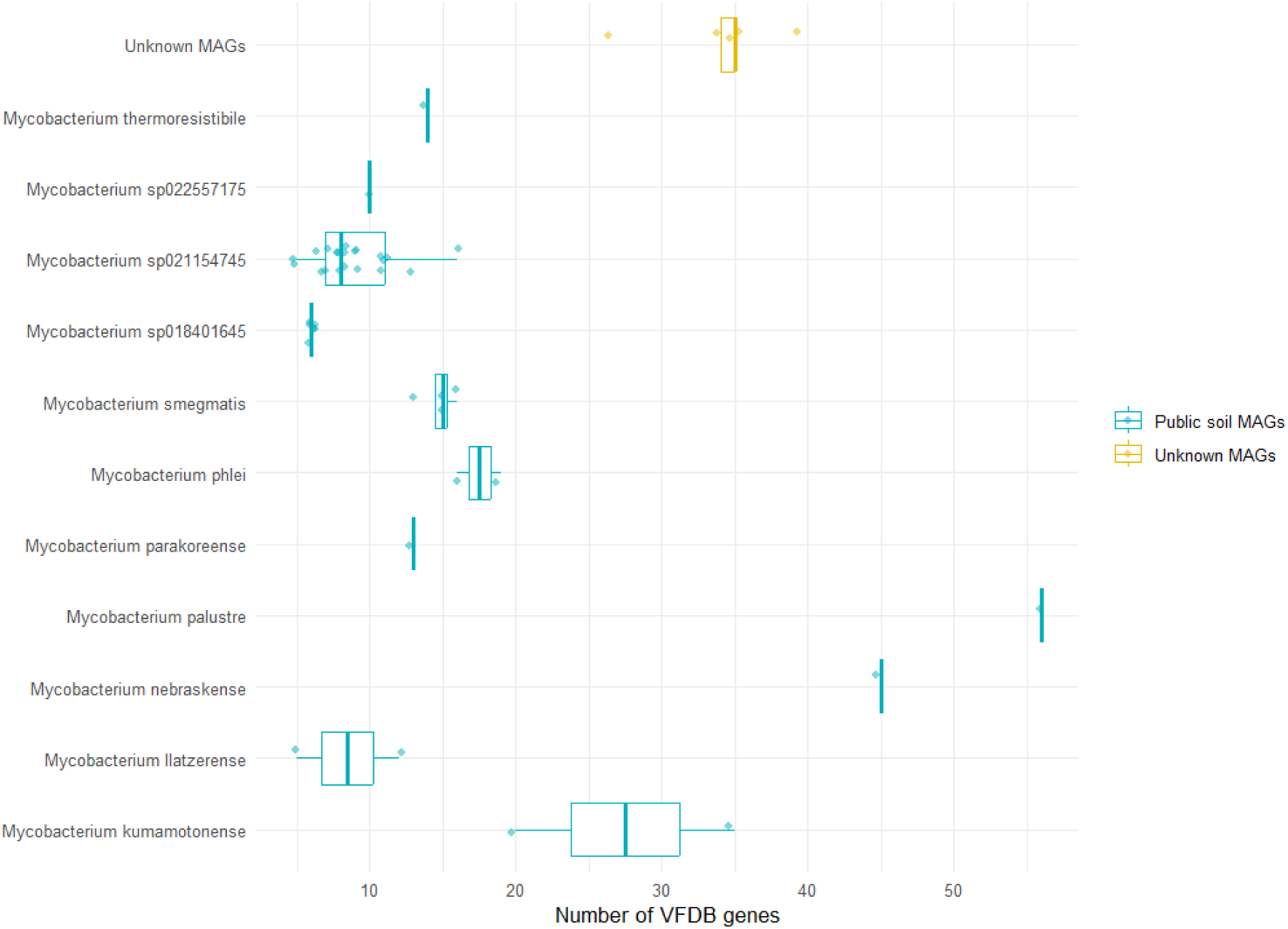
Comparison of virulence factor content within the VFDB of the unknown mycobacterium bins reconstructed here (Yellow) and mycobacterium MAGs reconstructed from soil samples elsewhere^24^ (Blue). The density of virulence factors is similar to other NTM MAGs like Mycobacterium kumamotonense^115^ and Mycobacterium phlei^116^. Data are represented as boxplots in which the middle line is the median, the lower and upper hinges correspond to the first and third quartiles, the upper whisker extends from the hinge to the highest value no further than 1.5 × interquartile range (IQR) from the hinge and the lower whisker extends from the hinge to the lowest value no further than 1.5 × IQR of the hinge.

**Figure S4.**
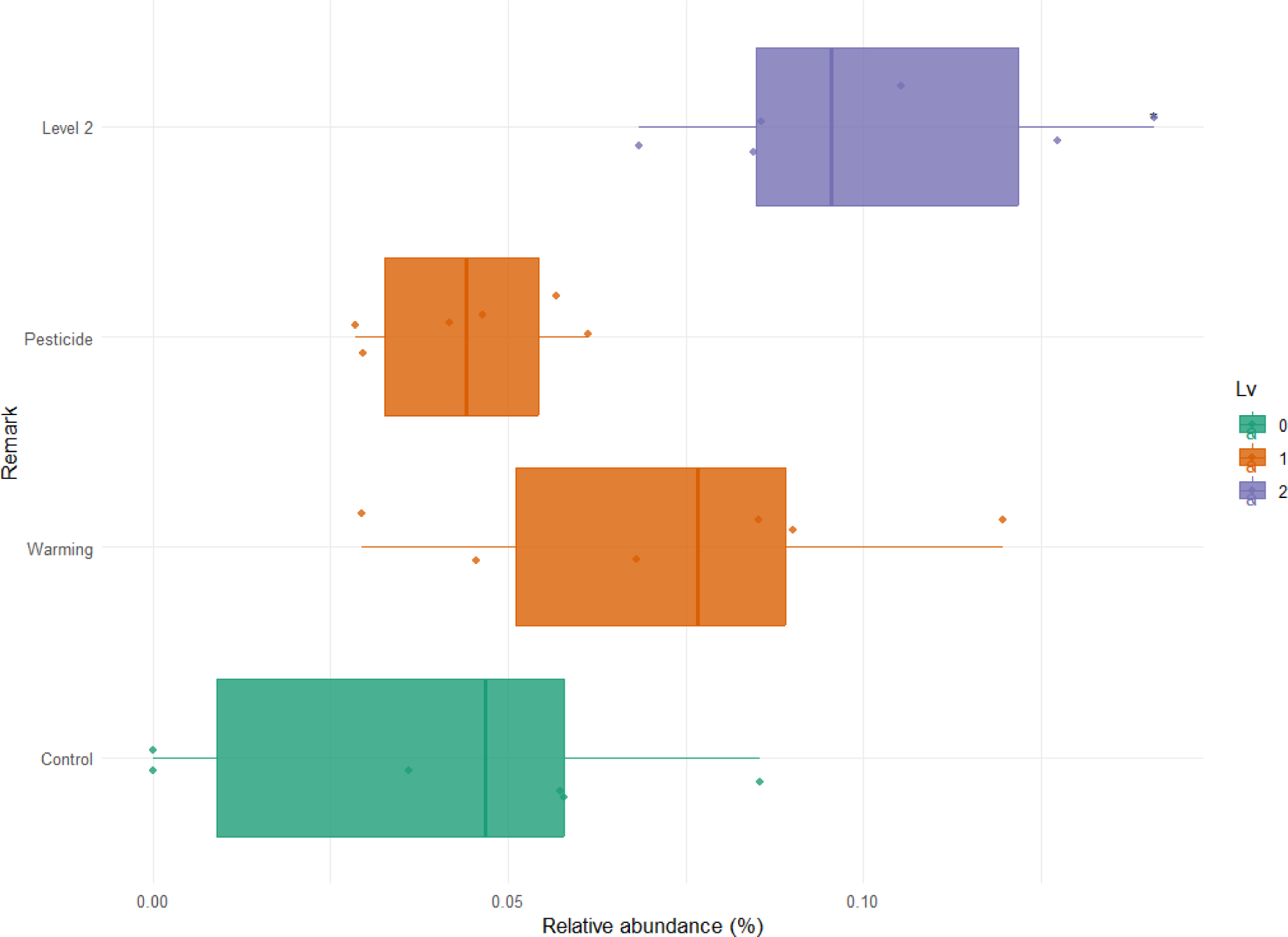
Relative abundance of the mycobacterium ASVs located in the freshwater stressor experiment in Romero, et al. (2020)^35^. Data are represented as boxplots in which the middle line is the median, the lower and upper hinges correspond to the first and third quartiles, the upper whisker extends from the hinge to the highest value no further than 1.5 × interquartile range (IQR) from the hinge and the lower whisker extends from the hinge to the lowest value no further than 1.5 × IQR of the hinge. Asterisks represent different significance levels obtained after a Two-sided Wilcoxon test with control samples; * indicate p-value <= 0.05, ** p-value <= 0.01, *** p-value <= 0.001 and **** p-value <= 0.0001

**Figure S5.**
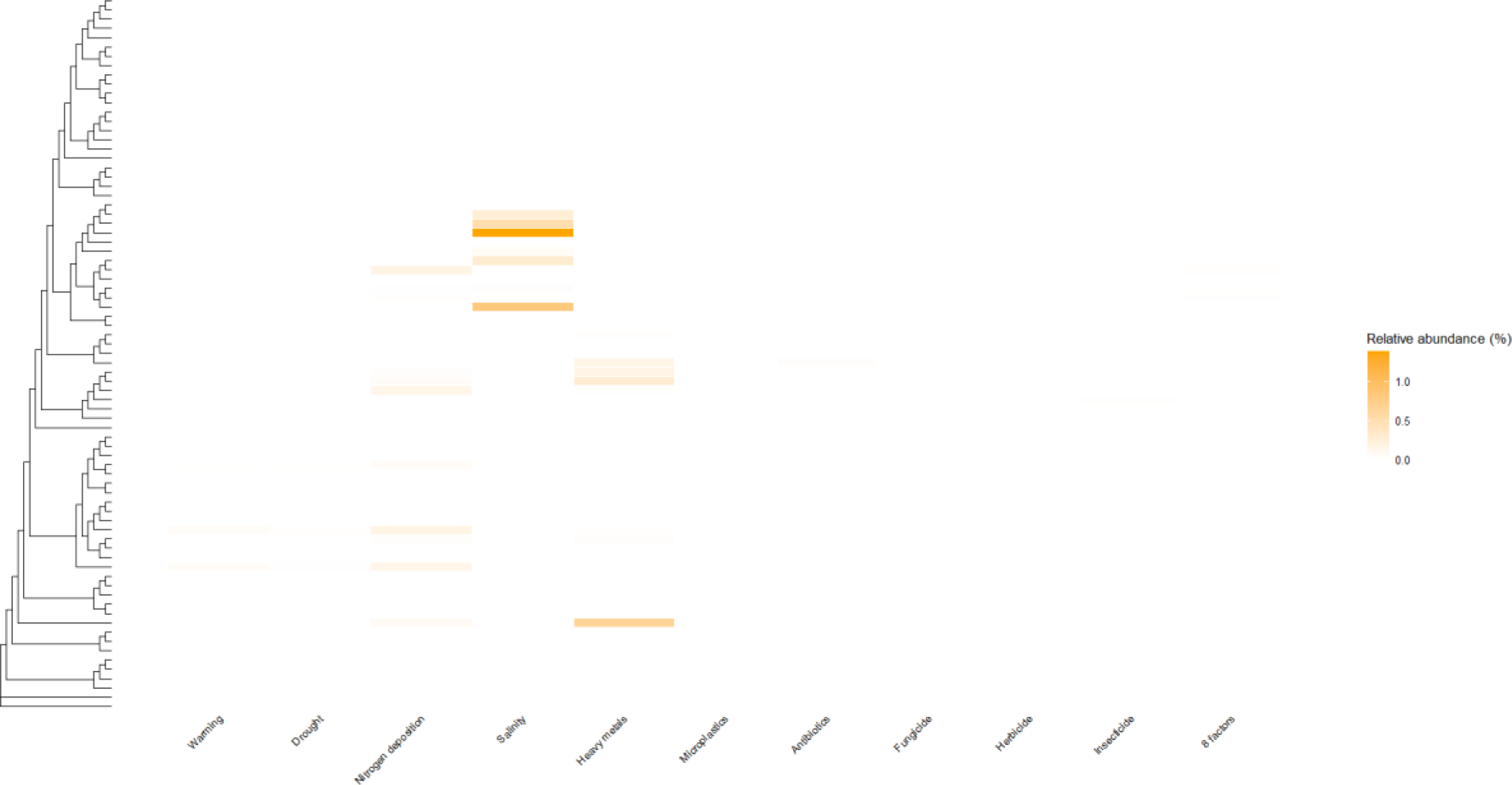
Mean relative abundance of MAGs not detected in most control samples (median abundance = 0) after the different GC treatments. For clarity, the abundance for MAGs with median relative abundance higher than 0 in the control samples were not included in the heatmap.

**Figure S6.**
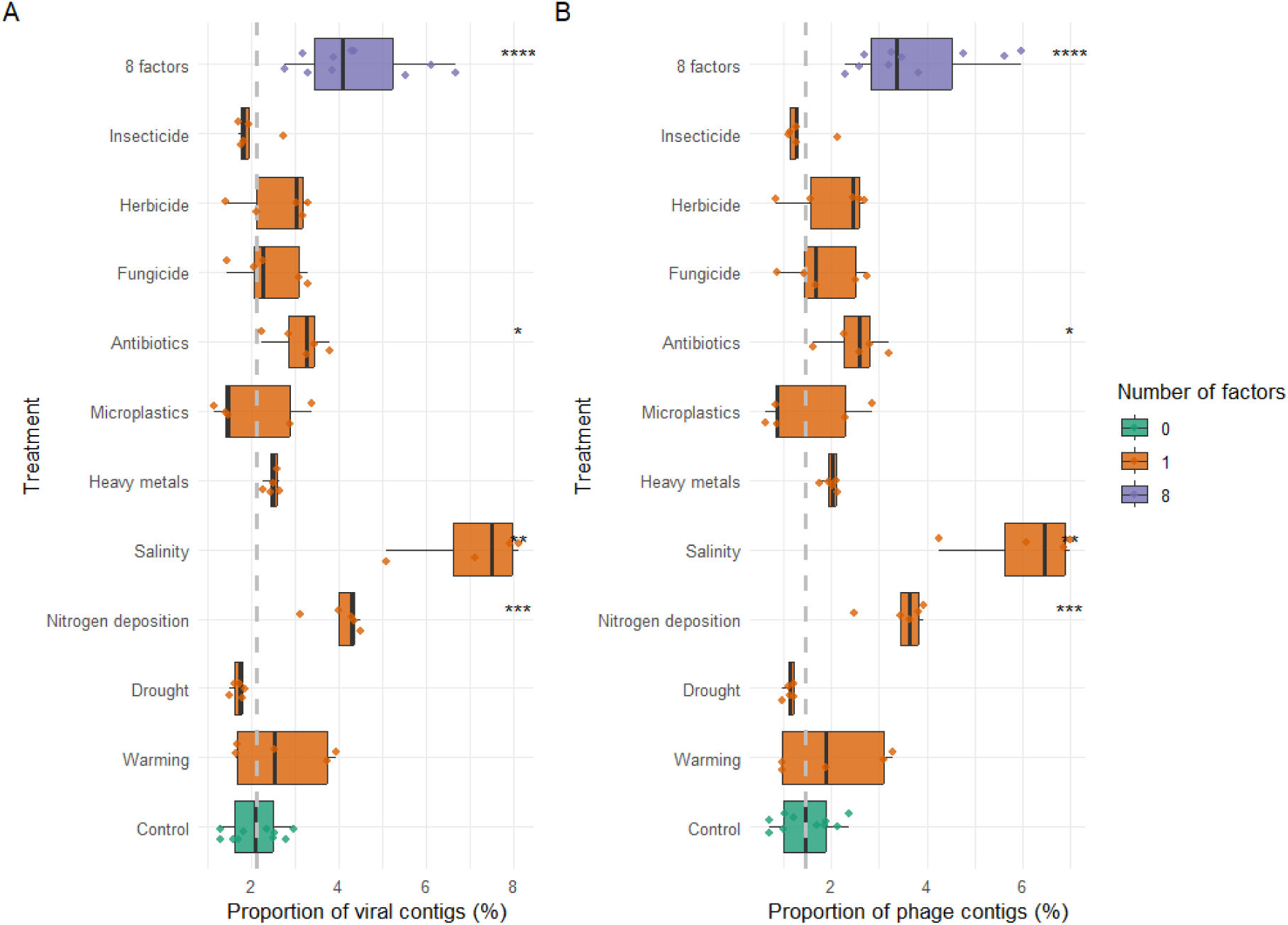
Proportion of A) viral contigs and B) phage contigs, per treatment. Data are represented as boxplots in which the middle line is the median, the lower and upper hinges correspond to the first and third quartiles, the upper whisker extends from the hinge to the highest value no further than 1.5 × interquartile range (IQR) from the hinge and the lower whisker extends from the hinge to the lowest value no further than 1.5 × IQR of the hinge. Asterisks represent different significance levels obtained after a Two-sided Wilcoxon test with control samples; * indicate p-value <= 0.05, ** p-value <= 0.01, *** p-value <= 0.001 and **** p-value <= 0.0001

**Figure S7.**
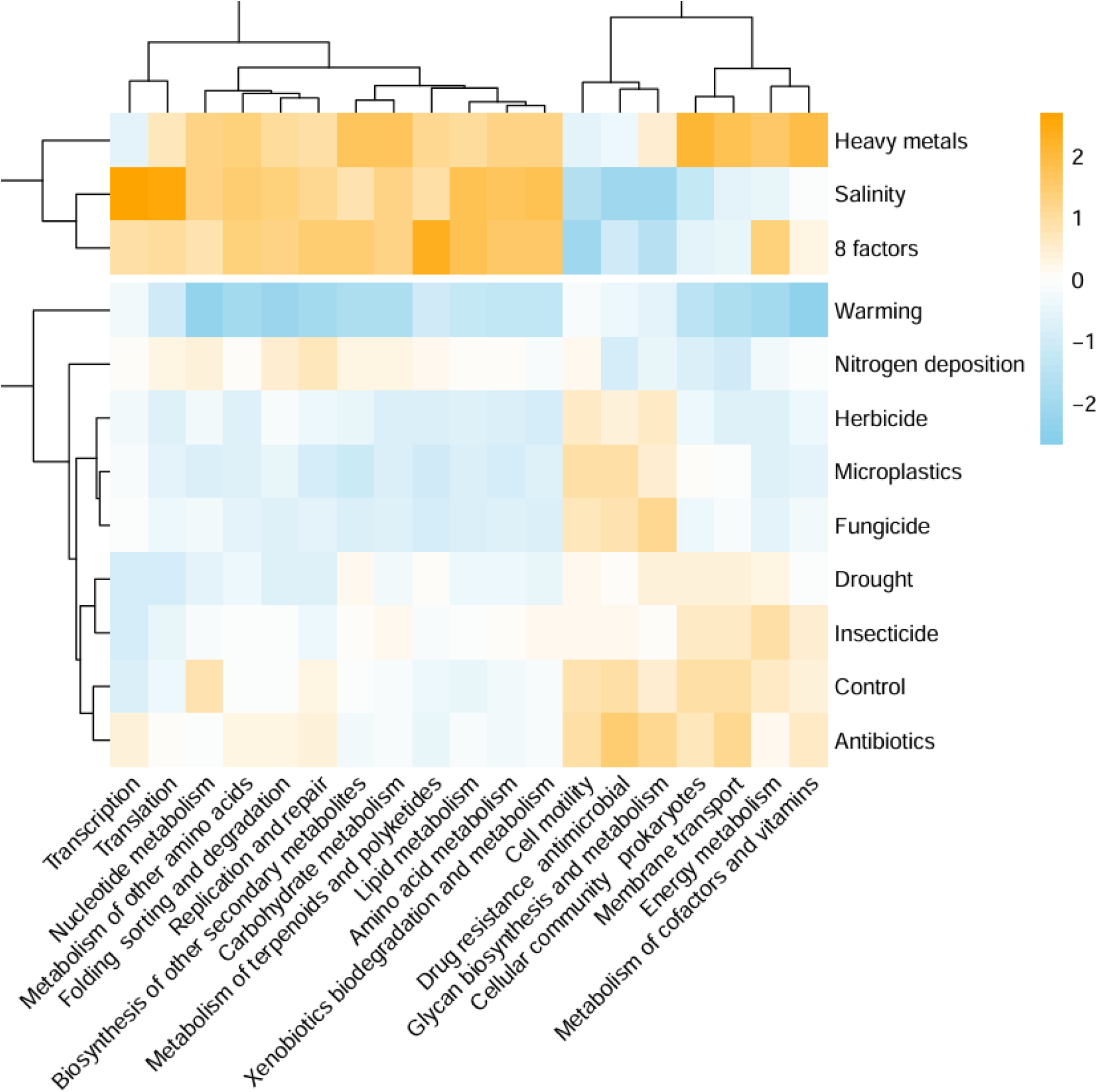
Heatmap indicating the copy number per cell, normalized by column, of general KEGG pathways.

**Figure S8.**
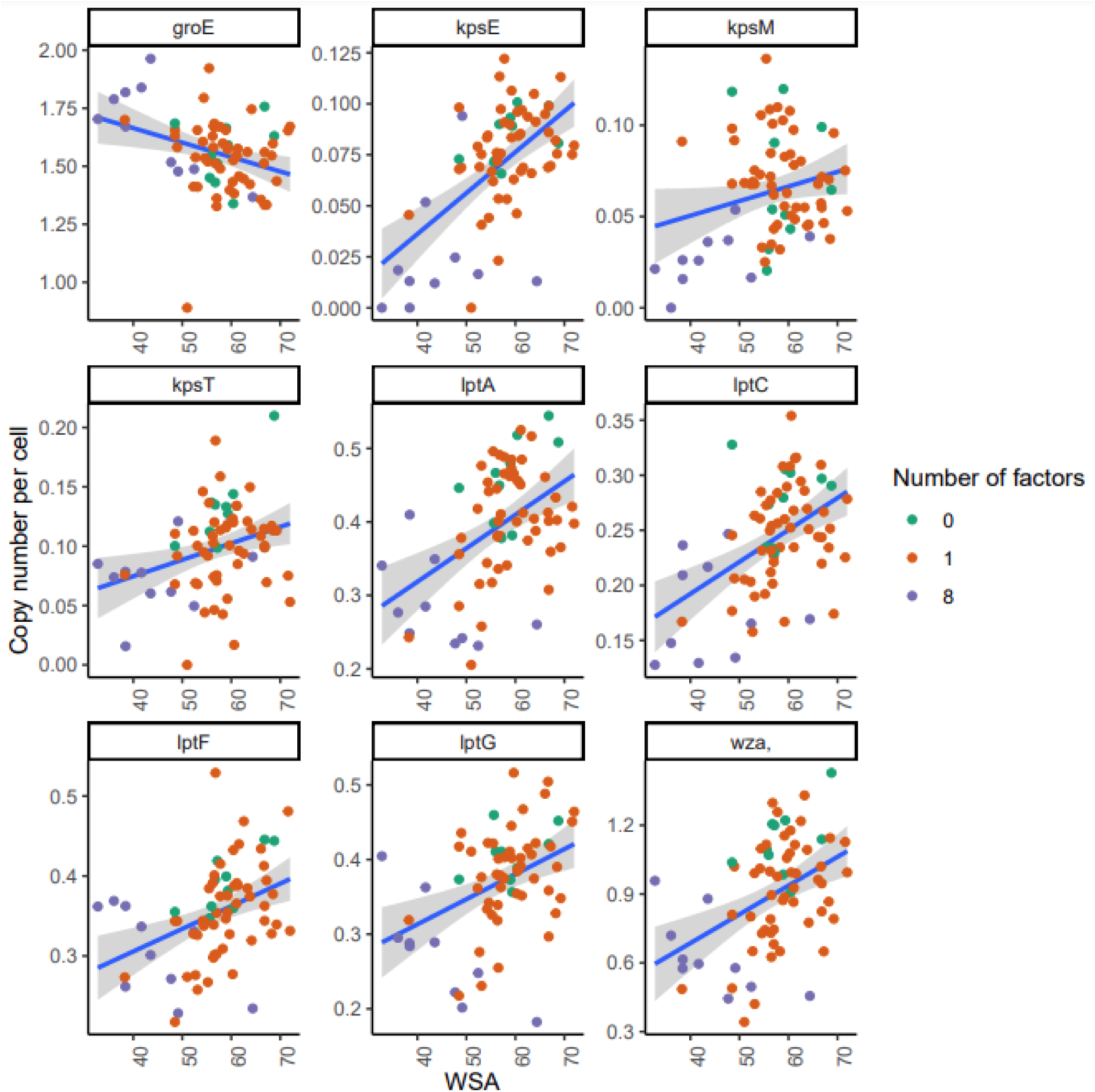
Correlation of genes previously associated with Water Stable Aggregates (WSA), with WSA measures.

**Figure S9.**
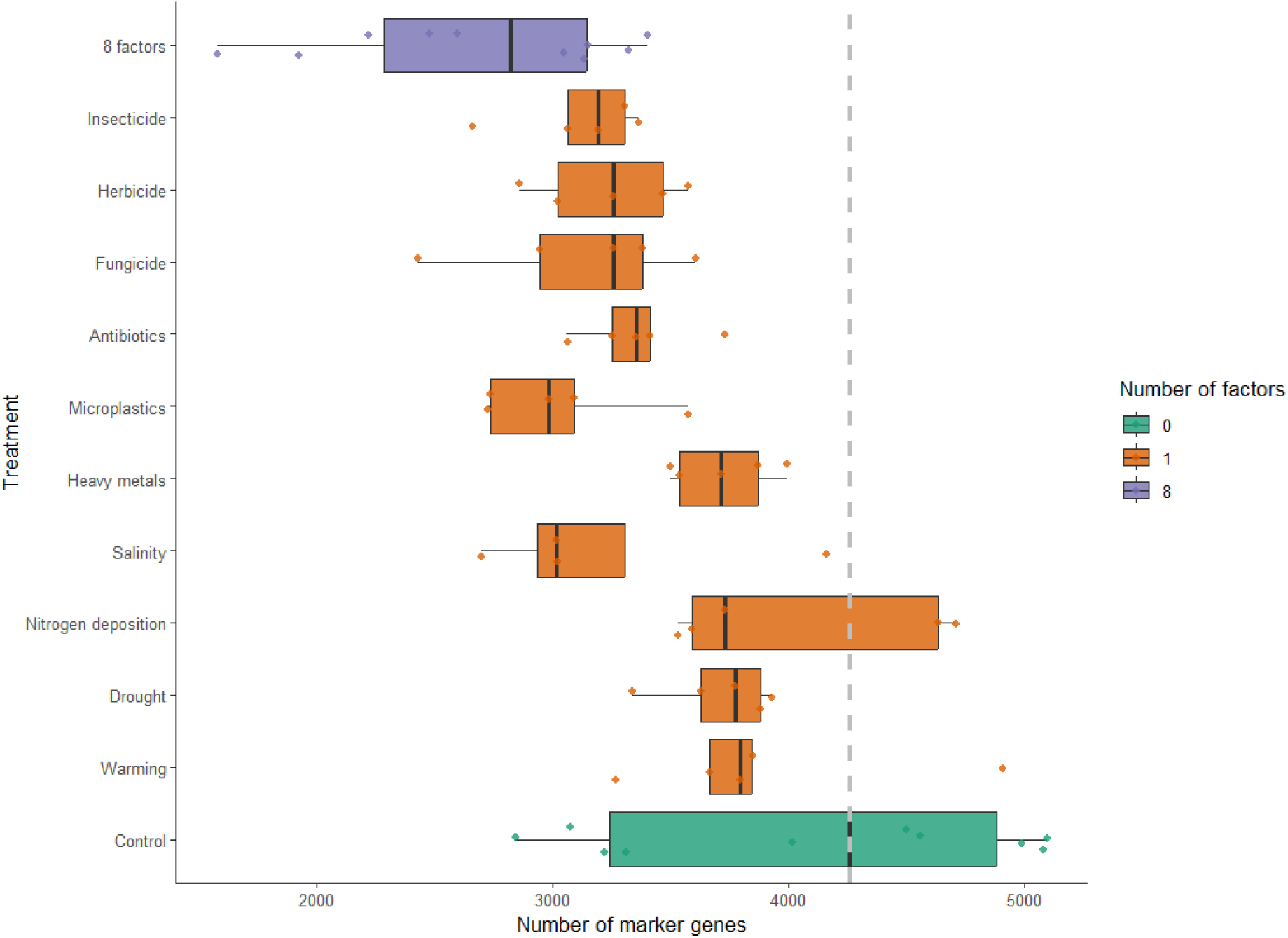
Number of marker genes, per treatment. Data are represented as boxplots in which the middle line is the median, the lower and upper hinges correspond to the first and third quartiles, the upper whisker extends from the hinge to the highest value no further than 1.5 × interquartile range (IQR) from the hinge and the lower whisker extends from the hinge to the lowest value no further than 1.5 × IQR of the hinge.

## Supplementary tables

Table S1. Number of High Quality (HQ) and Medium Quality (MQ) genomic bins obtained with different approaches

Table S2. Genes enriched in conditionally rare bins increasing in abundance after the copper treatment with q-value < 1e-20.

Table S3. Genes enriched in conditionally rare bins increasing in abundance after the copper treatment with q-value < 1e-2.

